# Functional Test of a Naturally Occurred Tumor Modifier Gene Provides Insights to Melanoma Development

**DOI:** 10.1101/2023.11.14.567049

**Authors:** Mateo Garcia-Olazabal, Mateus Contar Adolfi, Brigitta Wilde, Anita Hufnagel, Rupesh Paudel, Yuan Lu, Svenja Meierjohann, Gil G. Rosenthal, Manfred Schartl

## Abstract

Occurrence of degenerative interactions is thought to serve as a mechanism underlying hybrid unfitness. However, the molecular mechanisms underpinning the genetic interaction and how they contribute to overall hybrid incompatibilities are limited to only a handful of examples. A vertebrate model organism, *Xiphophorus*, is used to study hybrid dysfunction and it has been shown from this model that diseases, such as melanoma, can occur in certain interspecies hybrids. Melanoma development is due to hybrid inheritance of an oncogene, *xmrk*, and loss of a co-evolved tumor modifier. It was recently found that *adgre5*, a G protein-coupled receptor involved in cell adhesion, is a tumor regulator gene in naturally hybridizing *Xiphophorus* species *X. birchmanni* and *X. malinche*. We hypothesized that one of the two parental alleles of *adgre5* is involved in regulation of cell proliferation, migration and melanomagenesis. Accordingly, we assessed the function of *adgre5* alleles from each parental species of the melanoma-bearing hybrids using *in vitro* cell proliferation and migration assays. In addition, we expressed each *adgre5* allele with the *xmrk* oncogene in transgenic medaka. We found that cells transfected with the *X. birchmanni adgre5* exhibited decreased proliferation and migration compared to those with the *X. malinche* allele. Moreover, *X. birchmanni* allele of *adgre5* completely inhibited melanoma development in *xmrk* transgenic medaka, while *X. malinche adgre5* expression did not exhibit melanoma suppressive activity in medaka. These findings showed that *adgre5* is a natural melanoma suppressor and provide new insight in melanoma etiology.

## Introduction

Almost a century ago, it was discovered that certain hybrids of the southern platyfish (*X. maculatus*) and green swordtails (*X. hellerii*) develop highly malignant melanoma (C 1928, G. 1928) due to the inheritance of an oncogene, and conditional loss of a hypothetical tumor modulator (For review: see (Schartl and Walter 2016, Schartl and Lu 2024)). From this hybrid crossing experiment, a possible genetic explanation to how hybrid melanoma is generated is that certain individuals of the platyfish carry an oncogene controlled by a tumor suppressor. In this case, the oncogene effect on the platyfish is only a local dysplasia of melanocytes visible as a black pigment spot of macromelanophores. The oncogene and the tumor suppressor are located in different linkage groups, thus, when a platyfish is crossed with a swordtail that lacks both genes, some individuals will express the oncogene without the control of the tumor suppressor and therefore develop malignant melanoma.

It has been found that a mutant duplicate of *Xiphophorus* Epidermal Growth Factor Receptor (*egfrb*) named *Xiphophorus melanoma receptor kinase* (*xmrk*) is a bona fide oncogene driving the melanomagenesis observed in *Xiphophorus* hybrids, whose expression is necessary and sufficient for tumor development in *Xiphophorus* (Schartl M 1999, Schartl M 2010, Schartl and Lu 2024). The *xmrk* driven melanoma is thought to be a postzygotic mechanism for species isolation, and the *xmrk* gene is an example of the few speciation genes identified to date (M 2008, Flachs, Mihola et al. 2012). However, other than tumor modifiers identified from laboratory interspecies hybrids, there is no real-life example showing that *xmrk* is involved in species isolation until a study recently identified natural hybrid zones involving *X. birchmanni* and X. *malinche* (Kazianis, Gutbrod et al. 1998, Lu Y 2020, Powell D 2020). *X. birchmanni* has *xmrk* and is polymorphic for a black spot in their caudal fin, comprised of hyperplastic melanophores. *X. malinche* lacks *xmrk* as well as the caudal fin black spot. Interspecies hybrids between the two can develop melanoma. Powell et al. 2020 performed an admixture mapping study of these hybrids and revealed that melanoma occurs when the hybrid inherited *xmrk* from *X. birchmanni* and the *X. malinche* allele of *adgre5* (Powell D 2020). The *adgre5* gene encodes a G-protein couple receptor, with varied roles in cell adhesion and signaling (Yona S 2008). It plays a vital role in the adhesion and migration of healthy immune cells, however, it is also a mediator of invasion in a variety of human cancers [reviewed in (Safaee M 2013)).The *X. birchmanni* and *X. malinche adgre5* alleles are different by five amino acids, including one in a conserved epidermal growth factor–like calcium binding site (Powell D 2020), These evidences suggested that *X. birchmanni adgre5* is a *xmrk* regulator while the *X. malinche* allele is insufficient for this function.

Despite the finding that *adgre5* is associated to melanoma tumor suppression, there is no functional test to validate the activity of the *adgre5*. Therefore, this study aimed to test both *in vitro* and *in vivo* if the *X. birchmanni adgre5* can inhibit melanoma development. We hypothesize that *adgre5* can inhibit melanocyte proliferation and migration. To test this hypothesis, we expressed the *X. birchmanni* and *X. malinche adgre5* in mammalian melanocyte and assessed melanocyte proliferation and invasion. In addition, we introduced both *adgre5* alleles separately into *xmrk-*transgenic medaka that develop pigment cell lineage-specific tumors to evaluate if the two parental species alleles are different in arresting *xmrk* oncogenicity.

## Methods

### Cell culture and generations of stable adgre5 expression lines

Murine melanocyte Melan-A cells were cultured in DMEM with pyruvate (Gibco # 11995073), supplemented with 10% Fetal Calf Serum, 1% penicillin/streptomycin and maintained at 37°C, 5% CO_2_ with 100% humidity.

To generate doxycycline-inducible expression cell lines, the vector pSB-ET-iE (M. Gessler, Dept. of Developmental Biochemistry, University of Wurzburg) was used. It allows integration of genes by *sleeping beauty*-mediated transposition. Dox-inducible promoters enable precise regulation of the expression of the targeted gene, facilitating elegant experimental designs. Here, the responsive T6 promoter drives expression of *adgre5* and EGFP, with an IRES site between them (Figure S1a). After transfection, these cells were selected with 1 μg/ml puromycin for 2 weeks.

*X. malinche* and *X. birchmanni* alleles of *adgre5* were amplified using primers with XbaI and ClaI restriction enzyme sites. PCR amplification was performed using cDNA from tissue samples of organisms of *X. birchmanni* and *X. malinche*. The High Fidelity Q5 polymerase (New England Bio Labs #M0491S) was used for PCR amplification. The respective restriction enzymes were used to clone the PCR product into the vector, and wild-type Melan-a cells were transfected using Fugene transfection reagent (Promega Catalog # E5911). As a result, pSB-ET-iE_*X. malinche*_*adgre5*, and pSB-ET-iE_*X. birchmanni*_*adgre5* stable cell lines were generated. No significant differences were observed in the expression levels of *adgre5* between melan-a cell lines containing the different alleles (Figure S2, ANOVA p value > 0.05 for differences between *X. birchmanni* and *X. malinche* adgre5 alleles).

### RNA extraction, cDNA synthesis and qPCR

RNA extraction, cDNA synthesis, and qPCR were performed to measure the gene expression levels of *adgre5* in the Melan-A cell lines transfected with different *adgre5* alleles. Total RNA was extracted from freshly harvested cells or cell pellets that were stored at -80°C, using TRIzol reagent according to the manufacturer’s protocol. DNase I digestion was carried out for 1 hour at 37°C to eliminate any contaminating DNA. RNA concentration was determined using a NanoDrop spectrophotometer. RevertAid First Strand cDNA Synthesis Kit and random hexamer primers were used to reversely transcribe 100-4000 ng of RNA according to the manufacturer’s instructions (Thermo Fisher Catalog # K1622). Fluorescence-based RT-qPCR was performed using SYBR Green reagent and analyzed with a Mastercycler ep Realplex. Gene expression levels were normalized to a housekeeping gene (*Hprt* fwd: ACTGGCAACACTAACAGGACT *Hprt* rev: TGTTGTTGGATATGCCCTTG, *adgre5* fwd: CATCCGGCCCCTTTACTTGT *adgre5* rev: GGGCCAAAGAAGCTCCAGAT) using the delta-delta Ct method, and the success of transfection was confirmed by checking for GFP expression using an inverted fluorescence microscope with a 63x objective (Figure S1c). Three independent replicates of cells were used for each treatment and each cell line.

### Cell proliferation assay

In triplicate, cells were counted and seeded at equal density (1-2 × 10^3^ cells/well) in 96-well plates. Cells were treated with either dox 0 or dox 500 (500 mg/ml) by adding the respective dox concentrations to cell media. On days 2, 3, and 4 after treatment, 5 mg/ml of 3-(4,5-dimethylthiazol-2-yl)-2,5-diphenyltetrazolium bromide (MTT) was added to each well at a ratio of 1:5 (MTT:medium). The medium was aspirated after 2 hours of incubation at 37°C, and 150 μl of DMSO was added to each well. The plate was then incubated on a shaking device at room temperature for 15 minutes. A microplate reader was used to measure formazan accumulation by reading the optical density at 590 nm with a reference filter of 620 nm. Quintuples for each treatment, species, and duration of the experiments were performed. Cell growth was calculated by subtracting the optical density at 590 nm observed in dox 0 cells from the optical density at 590 nm observed in dox 500 induced cells. An ANOVA and a Tukey post hoc test were used to determine the statistical differences between *adgre5 X. birchmanni* and *X. malinche* alleles in cell growth.

### Trans-well migration assay

To assess cell migration, cells previously starved in 1% dialyzed FCS for 24 hours, were seeded at 2x10^3^ per well in the upper layer of uncoated trans-well inlays with 8 μm pore diameter in 24-well plates. The cells were treated with either dox 0 or dox 500 (500mg/ml) for 24 hours. To stimulate migration, medium containing 10% FCS and either dox 0 or dox 500 (500mg/ml) was added to the lower layer of the trans-well, and the cells were allowed to migrate for 16 hours.

Each assay was performed in triplicate per cell line and per dox treatment. Non-migrated cells were removed by cotton swabs, and the migrated cells were fixed with methanol, stained with 0.2% crystal violet dye in 2% ethanol for 15 minutes, and washed with PBS. The membrane was cut out of the trans-well inlay, embedded with Mowiol (polyvinyl alcohol) on microscope slides, and images were captured. The migrated cells were counted under a microscope, and migration was calculated as the difference in the number of cells that migrated in dox 500-induced cells minus the number of cells that migrated in dox 0 cells.

### Isolation of the *Xiphophorus adgre5* and construction of expression vectors with *fugu tyrp* promoter

The *adgre5* was designed to be driven by a pigment cell specific promoter, *Fugu rubripes* Tyrosinase-related Protein 1 gene (*tyrp*) (F 1996, Zou J 2006). To isolate the *adgre5* gene from *Xiphophorus*, a high-fidelity PCR (using Q5 Taq enzyme) was performed from cDNA extracted fin tissue from either *X. birchmanni* or *X. malinche* using overhang primers containing the XbaI restriction enzyme cutting site. Each cloning step was performed according to manufacturer’s protocols and plasmids were extracted using Quiagen Mini Prep Kit. Plasmids were sequenced to verify that no mutations occurred during cloning.

mCerulean was extracted with its promoter and a bGH poly(A) site using KpnI restriction enzyme from the brainbow vector. The pIsceI-typr-pa 3 plasmid containing the *tyrp fugu* promoter was linearized with KpnI restriction enzyme and ligated to ……. The resulting vector was subsequently digested with XbaI and ligated to the previously XbaI digested *adgre5* PCR products. Thus, specific plasmids were created containing either the *X. malinche* or *X. birchmanni* alleles of *adgre5* under the expression of the *tyrp fugu* promoter (Figure S3)

### Generation of transgenic medaka

Animal models are imperative to evaluate their importance in the context of the whole organism. Moreover, the advantage of establishing transgenic organisms is that we can study the isolated effects of the induced gene without other unknown possible genetic interactions. To generate stable transgenic lines, the meganuclease injection protocol was used since it was demonstrated to be more effective than injecting the plasmid alone, as it reduces mosaic expression, increases frequency of positive founder fish and increases germline transmission rates (Thermes et al., 2002). One-cell stage *tg*(*mitf*:*xmrk*) medaka embryos (*Oryzias latipes* strain: Carbio) were injected into the cytoplasm with approximately 15–20 pg of total DNA plasmid in a volume of 500 pl injection solution containing I-SceI meganuclease. Adult F0 fish were mated with each other and the offspring were tested for the presence of the transgene by screening of blue eyes under UV light (mCerulean effect).

All animal studies have been approved by the authors’ Institutional Review Board (Animal Welfare Officer of the University of Wurzburg). Adult fish were maintained under standard conditions (Kirchen and West, 1976) with an artificial photoperiod (10 hours of darkness, 14 hours of light) to induce reproductive activity. Clusters of fertilized eggs were collected 0.5–1 hour after the onset of light and kept in a rearing medium containing 0.1% NaCl, 0.003% KCl, 0.004% CaCl_2_ × 2H_2_O, 0.016% MgSO_4_ × 7H_2_O, and 0.0001% methylene blue.

F1 Fish (8 *tg*(*mitf*:*xmrk)+tyrp:Xmal*, and 7 *tg*(*mitf*:*xmrk)+tyrp:Xbir* were anesthetized in MS-222 and photographed with a Nikon D300 digital camera with a Tamron SP 90mm F/2.8 1:1 Macro lenses. Individuals were quantified for hyperpigmented melanic areas from the images using ImageJ (Schneider et al., 2012). Data fitted normality principles and therefore were analyzed with an ANOVA and a Tukey post hoc test.

## Results

### *X. malinche* but not *X. birchmanni* allele of *adgre5* promotes migration

To assess the potential role of *adgre5* in promoting cell migration, we performed tran-well migration assay of Melan-A cells expressing *X. birchmanni* and *X. malinche adgre5* respectively. Melan-A cells expressing the *X. birchmanni* allele of *adgre5* did not show difference in migration propensity compared to empty vector transfected cells. However, cells expressing the *X. malinche* allele of *adgre5* exhibited enhanced migration compared to both control cells and cells expressing *X. birchmanni adgre5* (Figure 1a, t-test p-value = 0.00582).

**Figure 1.**
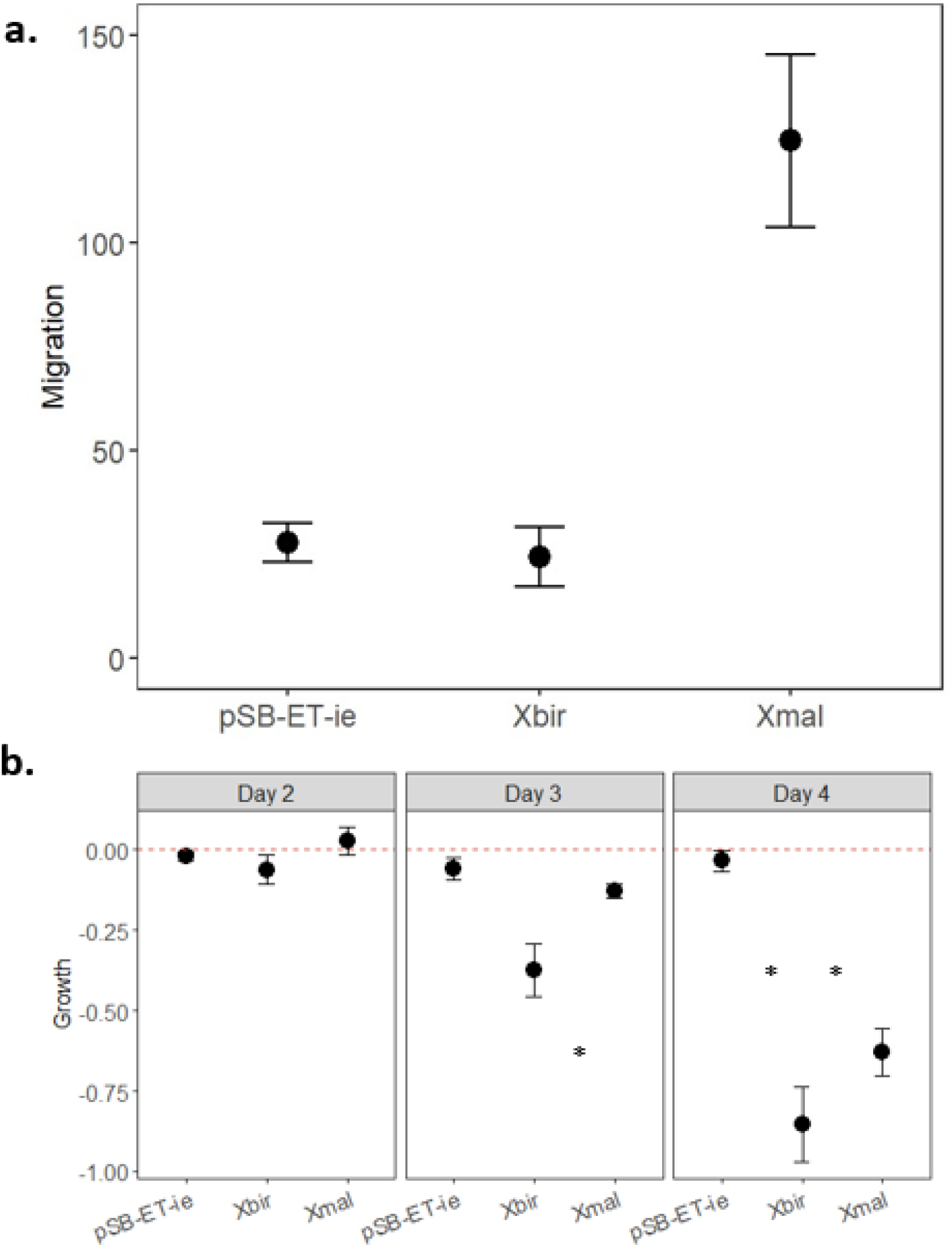
a) Migration assay (calculated as the difference between number of cells that migrated in dox 500 induced cells minus the number of cells that migrated in dox 0 cells). Xmal = stable cell line transfected with the *X. malinche* allele of *adgre5*. Xbir = stable cell line transfected with the *X. birchmanni* allele of *adgre5*.pSB-ET-ie….. b) Cell proliferation (calculated as the difference in optical density at 590 nm observed in dox 500 induced cells minus the optical density at 590 nm observed in dox 0 cells.) for each cell line and for each duration of the growth experiment. The dashed red line indicates no difference in growth between dox treated and untreated cells. The plots show the mean, and whiskers indicate two standard errors of the mean.

### *X. birchmanni* allele of *adgre5* inhibits pigment cell proliferation

Melan-A cells transfected with the *X. birchmanni* allele of *adgre5* showed lower cell numbers than control cells transfected with empty vector after 3 and 4 days of expression induction. In comparison, the cells expressing *X. malinche* adgre5 also exhibited lower cell number than control cells, but its effect is less potent than the *X. birchmanni* allele (Fig 1b, ANOVA p-value < 0.0001, Tukey post hoc p values, day 2: X.bir vs. X.mal p-value = 0.49; day 3: X.bir vs. X. mal p-value = 0.0006; day 4: X.bir-X.mal day 4 p-value= 0.0019).

### *X. birchmanni adgre5* eliminates *xmrk-*driven melanomagenesis in transgenic medaka

*Xiphophorus* is viviparous. Despite efforts are being put into development of producing transgenic *Xiphophorus* lines, there is currently no established protocol for this task. Therefore, we utilized an egg-laying fish model organism that is closely related to the *Xiphophorus*. The *xmrk—*transgenic medaka line develops pigment cell lineage-specific tumors with 100% genetic penetrance, and has been shown to be a valid model system to test gene functions for arresting the *xmrk* oncogenic activity (Patton and Nairn 2010, Schartl M 2010, Klotz, Kneitz et al. 2018, Regneri J 2019, Sugiyama, Schartl et al. 2019, Abdulsahib, Boswell et al. 2024). We expressed *X. birchmanni* and *X. malinche* alleles of *adgre5* in *xmrk-*transgenic medaka respectively [*tg(mift:xmrk)+tyrp:X*.*bir_adgre5*, and *tg(mift:xmrk)+tyrp:X*.*mal_adgre5*]. The *tg(mift:xmrk)+tyrp:X*.*bir_adgre5*, showed significant reductions of pigmentation compared to control *tg(mift:xmrk)* fish [Fig. 2; p-value < 0.00001], or *tg(mift:xmrk)+tyrp:X*.*mal_adgre5* medaka [Fig. 2; p-value < 0.00001], and devoid of pigment cell tumor. In comparison, the *tg(mift:xmrk)+tyrp:X*.*mal_adgre5* and *tg(mitf:xmrk)* medaka lines did not show significant differences in pigmentation to *xmrk* control medaka [Tukey post hoc p values, *tg(mift:xmrk)+tyrp:X*.*mal_adgre5*-*tg(mitf:xmrk)* = 0.85).

**Figure 2.**
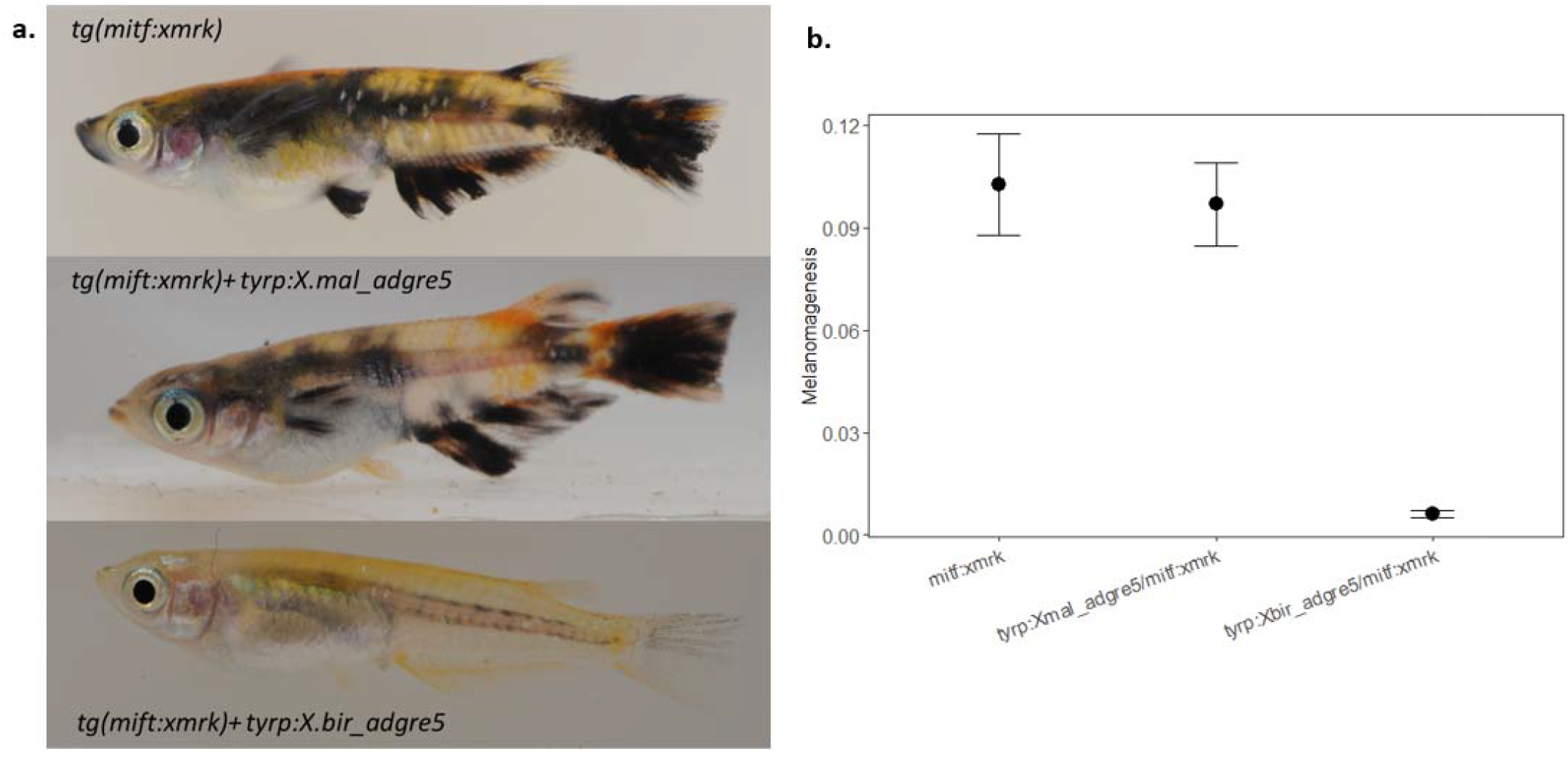
a) Phenotypes of the single and double transgenic fish. All individuals photographed of *tg(mift:xmrk)+tyrp:X*.*mal_adgre5, tg(mift:xmrk)+tyrp:X*.*bir_adgre5* are F0s, *tg(mitf:xmrk)* are more than 30 generations old. b) Melanomagenesis area (corrected by standard length) for each transgenic line generated. The plot shows the mean, and whiskers indicate two standard errors of the mean.

## Discussion

This study focuses on functionally testing the role of the *X. birchmanni* and *X. malinche adgre5* alleles by analyzing their effect on cell proliferation and migration and melanoma development. Melan-a cells are a well-established line of normal mouse melanocytes (Rossato F 2014) and provide an excellent non-tumorigenic line for studying the cellular and molecular basis of melanoma malignancy(Bennett D 1987), particularly for studies aimed at understanding the mechanisms triggering the change from benign pigmentation lesions to malignant melanoma. Two main characteristics of cancer are unlimited cell growth and potent metastatic properties (W 1956). Therefore, it is expected that tumor suppressor genes cause a decrease of proliferation and lowered propensity in migration. Even though melanoma modifying genes are easiest to investigate in cell culture models, animal models are imperative to evaluate their importance in the context of the whole organism. Transgenic organisms isolate effects of the induced gene from other unknown possible genetic interactions. The lack of differences in the areas covered by melanoma cells between the *tg(mift:xmrk)+tyrp:X*.*mal_adgre5* and *tg(mitf:xmrk)*, as well as the striking reduction of pigmentation of *tg(mift:xmrk)+tyrp:X*.*bir_adgre5*, suggest that *adgre5* acts indeed as a tumor suppressor. Moreover, pigment cells transfected with the *X. birchmanni* allele of *adgre5* caused cells in-vitro to grow slower and migrate less than those transfected with the *X. malinche* allele of *adgre5*.

*ADGRE5* is a prototypic member of the Adhesion class of G protein-coupled receptors (Adhesion GPCRs), which plays vital roles in numerous developmental processes as well as in tumorigenesis. Although it has been demonstrated that *adgre5*, under apoptotic conditions, can increase tumor cell viability by inhibition of caspase activation and modulation of anti- and pro-apoptotic members of the BCL-2 superfamily(Hsiao C 2015), it has also been demonstrated that G protein-coupled receptors inhibit melanoma tumor growth and metastasis (Xu L 2006). In fact, a recent review (T 2020) proposes Adhesion G protein–coupled receptors as candidate metabotropic mechanosensors and novel drug targets for numerous cancer types because they are surface molecules that act as mechanosensors. Information about *adgre5* function could also be expected from loss of function studies. Recently, it was demonstrated that *xmrk* can be suppressed when co-expressed with a known potent tumor suppressor in medaka transgenic lines: *cdkn2ab (Regneri J 2019)*. Future studies to further characterize the role of *adgre5* as a tumor suppressor are needed. For example, a tissue-specific knock-out of medaka adgre5 could cause even more malignant melanoma phenotypes, in analogy to the knockout of *cdkn2ab*.

The study by Powell et al. in 2020 was the first to propose a candidate hybrid incompatibility identified to the single-gene-level in naturally occurring hybrids. Almost simultaneously, another important example of hybrid incompatibility single-gene-level identification came from the closely related artificial hybrids between *X. maculatus* and *X. helleri (Powell D 2020)*. Lu et al., 2020 successfully identified *rab3d* as a tumor suppressor candidate gene. Future studies should focus on experimentally testing the role of *rabd3 (Lu Y 2020)* in a similar approach as taken here for *adgre5* using transgenic studies with the medaka *tg*(*mitf*:*xmrk*) line as well as cell culture experiments s.

The results of both studies are interesting because they propose that alleles of different genes, 7 Mb apart on the same chromosome, which interact with *xmrk*, are responsible for hybrid incompatibility in two different pairs of species from the same genus *Xiphophorus*. This suggests that a melanoma incompatibility involving *xmrk* originated independently in two distinct lineages. Schartl 2008 proposed that *R* (as it was called the then unknown tumor regulator gene) “could have preexisted before *xmrk* arose and would have suppressed the melanoma from the moment when the oncogene arose”. The most parsimonious hypothesis is that *xmrk* originated in a common ancestor to all *Xiphophorus* and that it has been repeatedly lost in several branches of the tree. In contrast, it appears that all species have *R*, but different alleles, because different hybrid crosses have different levels of *xmrk* suppression (J 1962, F 1967). Indeed, the recent findings suggest that R “different alleles” are actually different genes.

There has been renewed interest in the search for Bateson-Dobzhansky-Muller hybrid incompatibilities since we realized that hybridization is more common than previously thought (N 2002, Schumer M 2014). However, few studies have been able to precisely identify the interacting genes responsible for the incompatibility(D 2010). Of those that have, most identify candidate genes by using hybrids between crosses of model species that no longer hybridize naturally (Lee H 2008, Bikard D 2009, A 2018, Yu X 2018). The list of studies that have been able to pinpoint single gene effects grows significantly when considering which ones are actually able to experimentally test them and effectively assign a causal relationship between the gene interaction and the hybrid incompatibility(Trachtulec Z 2008, Mihola O 2009).

To the best of our knowledge, the work presented here is the only study so far that has experimentally validated a candidate gene involved in a hybrid incompatibility in species that currently hybridize naturally. Further studies of this kind could shed important light on how incompatibilities affect speciation, specifically whether mapped incompatibilities trigger divergence between species or arise after gene flow has ceased. The results of this study demonstrate that *adgre5* acts as a tumor modifier and highlight the importance of following population genetic mapping studies with functional cell culture and *in vivo* studies. Future research focused on characterizing the specific mechanism by which a*dgre5* suppresses *xmrk* holds great promise for biomedical research. In 2024, about 100,640 new melanomas will be diagnosed (about 59,170 in men and 41,470 in women, and about 8,290 people are expected to die of melanoma (about 5,430 men and 2,860 women) in the U.S. due to melanoma, which accounts for the vast majority of skin cancer deaths, despite only accounting for approximately 1% of all skin cancer cases (American Cancer Society, 2024). Insights into how a tumor suppressor gene can counteract an oncogene will be of tremendous value in the development of melanoma treatments.

## Acknowledgments

This work was funded by the Cancer Prevention and Research Institute of Texas, Grant ID RP200657, National Institutes of Health, National Cancer Institute award number R15 CA-223964, and Division of Comparative Medicine award number R24 OD-031467. MGO was funded by Fulbright and HIAS Scholar program

## CRediT author statement

**Mateo Garcia-Olazabal:** Conceptualization, Data curation, Formal analysis, Investigation, Project administration, Writing- original draft, Writing- review & editing **Mateus Contar Adolfi:** Conceptualization, Supervision, Writing- review & editing **Brigitta Wilde:** Investigation, Methodology **Anita Hufnagel:** Investigation, Methodology **Rupesh Paudel:** Investigation, Methodology, Writing- review & editing **Yuan Lu:** Conceptualization, Funding acquisition, Writing- review & editing **Svenja Meierjohann:** Conceptualization, Supervision, Writing – review & editing **Gil G. Rosenthal:** Funding acquisition, Supervision, Writing – review & editing **Manfred Schartl:** Conceptualization, Funding acquisition, Project administration, Supervision, Writing- review & editing.

## References

1-A, Z. M. a. S. (2018). “Gene duplicates cause hybrid lethality between sympatric species of Mimulus.” Plos Genetics 14: e1007130.

Abdulsahib, S., et al. (2024). “Transcriptional background effects on a tumor driver gene in different pigment cell types of medaka.” Journal of Experimental Zoology Part B: Molecular and Developmental Evolution 342(3): 252–259.

Bennett D C. P. a. H. I. (1987). “A line of non-tumorigenic mouse melanocytes, syngeneic with the B16 melanoma and requiring a tumour promoter for growth.” International Journal of Cancer 39: 414–418.

Bikard D P. D., Le-Metté C, Giorgi V, Camilleri C, Bennett M and Loudet O (2009). “Divergent Evolution of Duplicate Genes Leads to Genetic Incompatibilities Within A. thaliana.” Science 323: 623–626.

C, K. (1928). “Über Kreuzungen zwischen den Teleostiern Xiphophorus Helleri und Platypoecilus maculatus.” Zeitschrift für Induktive Abstammungs- und Vererbungslehre 47: 150–158.

D, P. (2010). “The molecular evolutionary basis of species formation.” Nature Review Genetics 11: 175–180.

F, A. (1967). “Tumour formation in Platyfish-Swordtail hybrids as a problem of gene regulation.” Experientia 23: 1–10.

F, d.-M. V. a. B. (1996). “Tyrosinase and related proteins in mammalian pigmentation.” FEBS Letters 381: 165–168.

Flachs, P., et al. (2012). “Interallelic and intergenic incompatibilities of the Prdm9 (Hst1) gene in mouse hybrid sterility.” Plos Genetics 8(11): e1003044.

G., H. (1928). “Uber Melanombildungen bei Bastarden von Xiphophorus maculatus var. rubra.” Klin Wochenschr 7: 1561–1562.

Hsiao C K. K., Chen H, Sittig D, Hamann J, Lin H and Aust G (2015). “The Adhesion GPCR CD97/ADGRE5 inhibits apoptosis.” The International Journal of Biochemistry & Cell Biology 65: 197–208.

J, A. (1962). “Effects of hybridization on pigmentation in fishes of the genus Xiphophorus.” Zoologica 47: 153–181.

Kazianis, S., et al. (1998). “Localization of a CDKN2 gene in linkage group V of Xiphophorus fishes defines it as a candidate for the DIFF tumor suppressor.” Genes, Chromosomes and Cancer 22(3): 210–220.

Klotz, B., et al. (2018). “Expression signatures of early-stage and advanced medaka melanomas.” Comparative Biochemistry and Physiology Part C: Toxicology & Pharmacology 208: 20–28.

Lee H C. J., Cheong L, Chang N, Yang S and Leu J (2008). “Incompatibility of nuclear and mitochondrial genomes causes hybrid sterility between two yeast species.” Cell 135: 1065–1073.

Lu Y S. A., Voss S, Lai Z, Kneitz S, Boswell W, Boswell M, Savage M, Walter C, Warren W, Schartl M and Walter R (2020). “Oncogenic allelic interaction in Xiphophorus highlights hybrid incompatibility.” Proceedings of the National Academy of Sciences of the USA 47: 29786–29794.

M, S. (2008). “Evolution of Xmrk: an oncogene, but also a speciation gene?” BioEssays 30: 822–832.

Mihola O T. Z., Vlcek C, Schimenti J and Forejt J (2009). “A Mouse Speciation Gene Encodes a Meiotic Histone H3 Methyltransferase.” Science 323: 373–375.

N, J. (2002). “Sixty years after “Isolating Mechanisms, Evolution and Temperature”: Muller’s legacy.” Genetics 161(3): 939–944.

Patton, E. E. and R. S. Nairn (2010). “Xmrk in medaka: A new genetic melanoma model.” Journal of Investigative Dermatology 130(1): 14–17.

Powell D G.-O. M., Keegan M, Reilly P, Du K, Díaz-Loyo A, Banerjee S, Blakkan D, Reich D, Andolfatto P, Rosenthal G, Schartl M and Schumer M (2020). “Natural hybridization reveals incompatible alleles that cause melanoma in swordtail fish “ Science 368: 731–736.

Regneri J K. B., Wilde B, Kottler V, Hausmann M, Kneitz S, Regensburger M, Maurus K, Götz R, Lu Y, Walter R, Herpin A and Schartl M (2019). “Analysis of the putative tumor suppressor gene cdkn2ab in pigment cells and melanoma of Xiphophorus and medaka.” Pigment Cell & Melanoma 32: 248–258.

Rossato F Z. K., La-Guardia P, Ortega R, Alberici L, Costa R, Catharino R, Graner E, Castilho R and Vercesi A (2014). “Fatty Acid Synthase Inhibitors Induce Apoptosis in Non-Tumorigenic Melan-A Cells Associated with Inhibition of Mitochondrial Respiration.” Plos One 9: e101060.

Safaee M C. A., Ivan M, Oh M, Bloch O, Sun M, Oh T and Parsa A (2013). “CD97 is a multifunctional leukocyte receptor with distinct roles in human cancers “ International Journal of Oncology 43: 1343–1350.

Schartl M H. U., Gutbrod H, Volff J and Wittbrodt J (1999). “Melanoma loss-of-function mutants in Xiphophorus caused by Xmrk-oncogene deletion and gene disruption by a transposable element.” Genetics 153: 1385–1394.

Schartl, M. and Y. Lu (2024). “Validity of Xiphophorus fish as models for human disease.” Disease Models & Mechanisms 17(1).

Schartl, M. and R. B. Walter (2016). “Xiphophorus and medaka cancer models.” Cancer and Zebrafish: Mechanisms, Techniques, and Models: 531–552.

Schartl M W. B., Laisney J, Taniguchi Y, Takeda S and Meierjohann S (2010). “A Mutated EGFR Is Sufficient to Induce Malignant Melanoma with Genetic Background-Dependent Histopathologies.” Journal of Investigative Dermatology 130(1): 249–258.

Schumer M C. R., Powell D, Dresner R, Rosenthal G and Andolfatto P (2014). “High-resolution mapping reveals hundreds of genetic incompatibilities in hybridizing fish species.” Elife e02535.

Sugiyama, A., et al. (2019). “Histopathologic features of melanocytic tumors in Xiphophorus melanoma receptor kinase (xmrk)-transgenic medaka (Oryzias latipes).” Journal of toxicologic pathology 32(2): 111–117.

T, L. (2020). “Adhesion G protein–coupled receptors—Candidate metabotropic mechanosensors and novel drug targets.” Basic & Clinical Pharmacology & Toxicology 126: 5–16.

Trachtulec Z V. C., Mihola O, Gregorova S, Fotopulosova V and Forejt J (2008). “Fine haplotype structure of a chromosome 17 region in the laboratory and wild mouse.” Genetics 178: 1777–1784.

W, O. (1956). “On the origin of cancer cells.” Science 123: 309–314.

Xu L B. S., Hearn J and Hynes R (2006). “GPR56, an atypical G protein_Jcoupled receptor, binds tissue transglutaminase, TG2, and inhibits melanoma tumor growth and metastasis.” Proceedings of the National Academy of Sciences of the USA 103: 9023–9028.

Yona S L. H., Siu W, Gordon S and Stacey M (2008). “Adhesion-GPCRs: emerging roles for novel receptors.” Trends in Biochemical Sciences 33: 491–500.

Yu X Z. Z., Zheng X, Zhou J, Kong W, Wang P, Bai W, Zheng H, Zhang H, Li J, Liu J, Wang Q, Zhang L, Liu K, Yu Y, Guo X, Wang J, Lin Q, Wu F, Ren Y, Zhu S, Zhang X, Cheng Z, Lei C, Liu S, Liu X, Tian Y, Jiang L, Ge S, Wu C, Tao D, Wang H and Wan J (2018). “A selfish genetic element confers non-Mendelian inheritance in rice.” 360: 1130–1132.

Zou J B. F., Wang J, Kawakami K and Wei X (2006). “The Fugu tyrp1 promoter directs specific GFP expression in zebrafish: tools to study the RPE and the neural crest-derived melanophores.” Pigment Cell Res. 19(6): 615–627.

